# Utilisation of the Prestwick Chemical Library ^®^ to identify drugs that inhibit the growth of Mycobacteria

**DOI:** 10.1101/357897

**Authors:** Panchali Kanvatirth, Rose E. Jeeves, Joanna Bacon, Gurdyal S. Besra, Luke J. Alderwick

## Abstract

Tuberculosis (TB) is an infectious bacterial disease that kills approximately 1.3 million people every year. Despite global efforts to reduce both the incidence and mortality associated with TB, the emergence of drug resistant strains has slowed any progress made towards combating the spread of this deadly disease. The current TB drug regimen is inadequate, takes months to complete and poses significant challenges when administering to patients suffering from drug resistant TB. New treatments that are faster, simpler and more affordable are urgently required. Arguably, a good strategy to discover new drugs is to start with an old drug. Here, we have screened a library of 1200 FDA approved drugs from the Prestwick Chemical library ^®^ using a GFP microplate assay. Drugs were screened against GFP expressing strains of *Mycobacterium smegmatis* and *Mycobacterium bovis* BCG as surrogates for *Mycobacterium tuberculosis,* the causative agent of TB in humans. We identified several classes of drugs that displayed antimycobacterial activity against both *M. smegmatis* and *M. bovis* BCG, however each organism also displayed some selectivity towards certain drug classes. Variant analysis of whole genomes sequenced for resistant mutants raised to florfenicol, vanoxerine and pentamidine highlight new pathways that could be exploited in drug repurposing programmes.

## Introduction

Tuberculosis (TB) remains a major global health issue, despite it being over twenty years since the World Health Organisation (WHO) declared TB a global emergency (1). In 2016, TB killed around 1.3 million people and now ranks alongside HIV as the leading cause of death globally. It has been estimated that almost 6.3 million new cases of TB are to have occurred in 2016; 46% of these new TB cases were individuals co-infected with HIV. Alarmingly, an estimated 4.1% of new TB cases and 19% of previously treated TB cases are infections caused by Multi-Drug Resistant TB (MDR-TB), and in 2016 an estimated 190,000 people died from this form of the disease. Furthermore, extensively drug-resistant TB (XDR-TB) has now been reported in 105 countries, and accounts for approximately 30,000 TB patients in 2016. If these numbers are to reduce in line with milestones set by the WHO End TB Strategy, alternative therapeutic agents that target novel pathways are urgently required.

Drug repurposing (or drug redeployment), is an attractive approach for the rapid discovery and, in particular, development of new anti-TB drugs (2, 3). Due to the time and cost of bringing new molecular entities through the developmental pipeline to clinic, drug repurposing offers an expedient option, in part due to pre-existing pharmacological and toxicological datasets that allow for rapid profiling of active hits (4). In this study, we used GFP-expressing strains of *M. smegmatis* and *M. bovis* BCG in order to screen the Prestwick Chemical Library ^®^ for antimycobacterial drugs. Together with drugs that have previously been identified from similar screens (5), we identified a number of novel hits that display good antimycobacterial activity which were also confirmed in*Mycobacterium tuberculosis* H37Rv. We sought to characterise the mode of action of selection of hits, by performing whole genome sequencing with variant analysis on laboratory resistant mutants. This study highlights both the usefulness and circumspection required when utilising *M. smegmatis* and *M. bovis* BCG in drug repurposing screens to new anti-TB agents.

## Results

### Primary screening of the Prestwick Chemical Library ^®^ against *M.smegmatis* and *M. bovis* BCG

To identify which of the 1200 FDA approved drugs in the Prestwick Chemical Library ^®^ inhibit the growth of mycobacteria, a high throughput fluorescence screen was used to measure GFP expression in strains of both *M. smegmatis* and *M. bovis* BCG (GFP microplate assay [GFPMA]) (6). *M. smegmatis*_pSMT3_eGFP and *M. bovis* BCG_pSMT3_eGFP were cultured in 96-well plates in the presence of 20 µM compound from the Prestwick Chemical Library ^®^. GFP fluorescence was measured at specified time points and data was normalized against both positive and negative controls to produce a scatter graph of the survival percentages (Fig. 1). In order to assess the reproducibility and robustness of the GFPMA HTS, we calculated Z’ factor values for each of the assay plates used to screen the 1200 compounds of the Prestwick Chemical Library ^®^against both *M. smegmatis*_pSMT3_eGFP and *M. bovis* BCG_pSMT3_eGFP. The Z’ values of the primary screen against *M. smegmatis* vary between 0.4 and 0.9 across each of the assay plates used in the screen (Fig. S1). In case of the primary screen for *M. bovis* BCG, the Z’values are consistent between 0.8 and 0.9 across all assay plates (Fig. S1). However, since all assay plates used in the experiment derived Z’ values ≥ 0.4, all data generated was deemed suitable for further processing (7). In order to understand the variability in the data and significance of the hits identified from the scatter plot (Fig. 1), we analysed the variance of data both between and across replicate experiments carried out using *M. smegmatis* and *M. bovis* BCG. The overall coefficient of correlation (r^2^) values for replicate assays were calculated to be 0.63 and to 0.89 for *M. smegmatis*_pSMT3_eGFP and *M. bovis* BCG_pSMT3_eGFP, respectively (Fig. 2). This indicates increased variance in the data for assays conducted with *M. smegmatis* compared to screens performed using *M. bovis* BCG. We analysed the frequency distribution of data both within and across each primary screen using *M. smegmatis* and *M. bovis* BCG (Fig. 3). For *M. smegmatis*, we observed that 31.5 % of the compounds screened in the library induced ≤ 75 % survival of bacterial cell growth in the primary screen (Fig. 3). For BCG, 21 % of the library induced ≤ 75 % survival of bacterial cell growth in the primary screen (Fig. 3). We applied a minimum cut-off of ≤25 % bacterial cell survival at 20 µM compound, as a parameter that defined an antimycobacterial hit that would be further investigated in downstream experiments (Fig. 1). In this regard, we observed an almost identical hit rate of 6.9% and 6.8% for compounds inducing ≤ 25 % survival for *M. smegmatis* and BCG, respectively (Fig. 3).

**Fig. 1.**
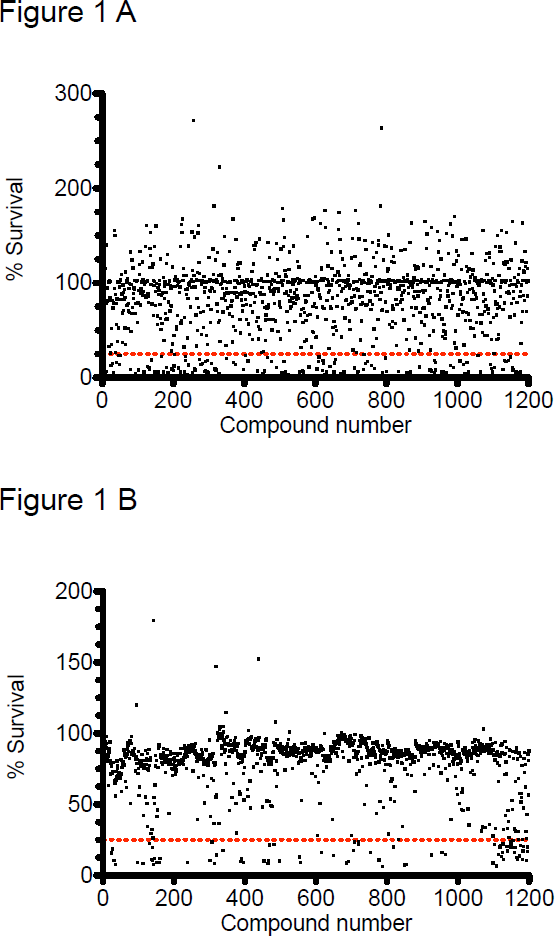
Primary screening of the Prestwick Chemical Library ^®^ compounds against*M. smegmatis* (A) and *M. bovis* BCG (B) using a GFPMA assay. GFP measurements were recorded after a defined period of incubation of mycobacteria in the presence of 20 µM compound from the Prestwick Chemical Library ^®^. Data was normalised to control wells and is expressed as mean % survival from n=2 biological replicate experiments. The red dashed line depicts <25 % cell survival as determined by residual GFP fluorescence.

**Fig. 2.**
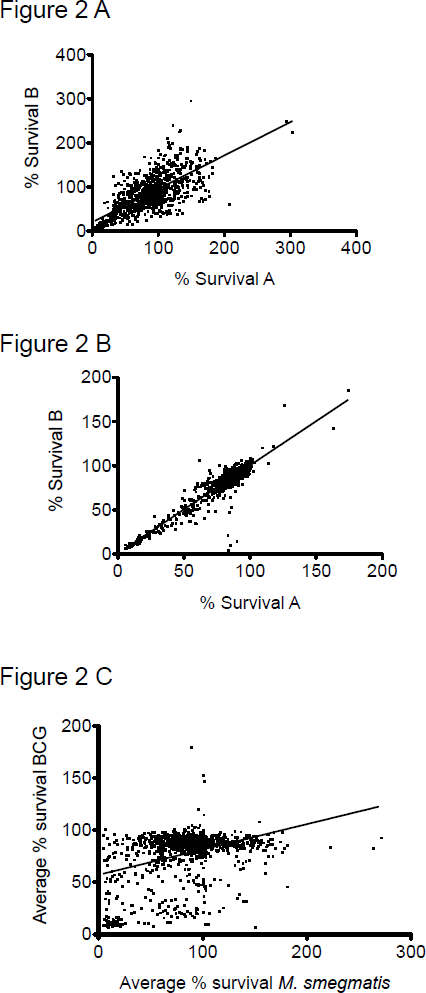
**Correlation analysis of the primary screen against the** Prestwick Chemical Library ^®^. Scatter graphs representing correlation analysis of the cumulative data of the percentage survivals between n=2 biological replicate experiments (run A and B) during the primary screen of the Prestwick Chemical Library ^®^ against *M. smegmatis* (A) and *M. bovis* BCG (B). The average of run A and B data sets from both *M. smegmatis* (A) and *M. bovis* BCG were plotted against each other (C).

**Fig. 3.**
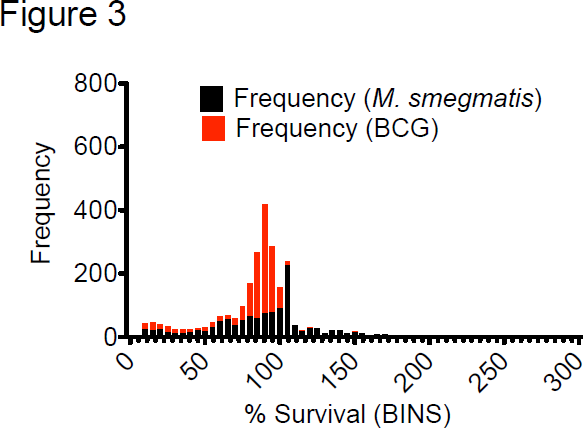
**A comparative frequency distribution of the primary screening against the** Prestwick Chemical Library ^®^. Each bar represents the comparative frequency distribution primary screening data (averaged) for *M. smegmatis* (black) and *M. bovis* BCG (red). % survival data for was ‘binned’ into groups of 5%.

The initial screen against *M. smegmatis* generated 83 hits which inhibited the survival of this fast-growing species of mycobacterium below 25% (Fig. 1A). The screen against the slower growing mycobacterial strain *M. bovis* BCG revealed 81 hits (Fig. 1A) which inhibited the growth of the bacteria below 25% (Fig. 1B). Categorisation of these hits (<25% survival) into pharmacological groups reveals that an almost equal number of fluoroquinolones, macrolides, polyketide antibiotics, antimycobacterial drugs, and antiseptics display inhibitory activity against both *M. smegmatis* and BCG whilst the aminoglycosides displayed more inhibitory activity towards *M. smegmatis* compared to BCG (Fig. 4). Other notable classes of drugs that inhibit the growth of both *M. smegmatis* and BCG include the amphenicols, glycopeptides and non-ribosomal peptide antibiotics, antihistamines, acetylcholine esterase inhibitors, antiemetic, antimalarial, antiprotozoal and surfactants (Fig. 4). Notable species-specific inhibitors affecting only *M. smegmatis* were also identified belonging to antiestrogen, antiarrhythmic and antipsychotic drugs (Fig. 4). For BCG, it appears that the cephalosporin antibiotics are only able to inhibit the slower growing mycobacterial species and do not affect the faster growing saprophytic organism *M. smegmatis.* Other significant drug classes that only inhibit BCG include anticancer agents, antidiabetics, anticonvulsants and angiotensin antagonists (Fig. 4).

**Fig. 4.**
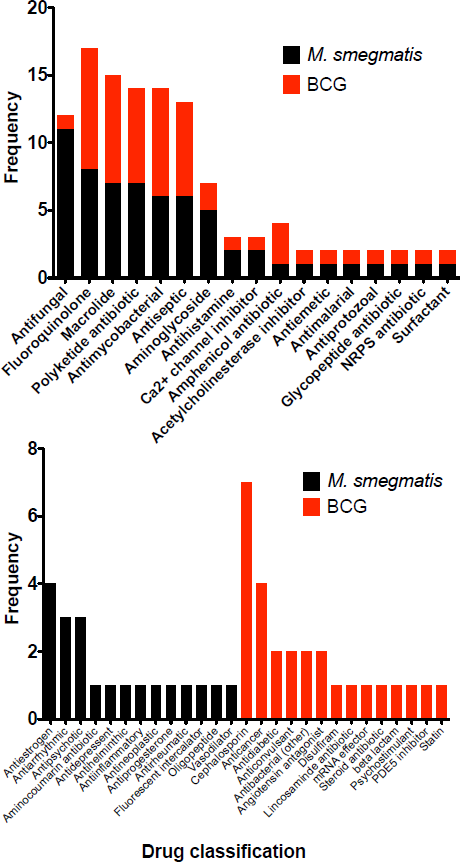
Comparison of the hits emerging from the primary screen active against both *M. smegmatis* and *M. bovis* BCG. Drugs were grouped into drug classifications and are plotted as frequency of hits against either *M. smegmatis* or *M. bovis* BCG.

All hits from the primary screen (displaying <25% survival) were filtered to remove all known antimycobacterial drugs and a significant number of other antimicrobial agents (5). The remaining drugs were then clustered into one of three groups based upon their inhibitory activity against either fast-growing (*M. smegmatis*) or slow-growing (*M. bovis* BCG) strains of mycobacteria, or those showing overlapping activity. Each cluster of drugs was further ranked and given a priority score that was based on the apparent potency of the drug and potential novelty of its mode of action from literature-based searches.

### Secondary screening and hit confirmation against *M.smegmatis*

Minimal Inhibitory Concentrations (MIC) were determined using the standardized broth dilution method and then subsequently measured on solid medium in order to ascertain the concentration required to generate resistant mutants (Fig. S2). Among the drugs tested against *M. smegmatis,* the most potent was meclocycline sulfosalicylate with a liquid and solid MIC of 0.10 µM and 0.2 µM, respectively (Table 1). Auranofin also displayed a relatively low solid MIC of 6.0 µM although this was 22-fold higher than its liquid MIC values (0.27 µM). Other drugs tested for MIC determination were alexidine and chlorhexidine which both exhibited relatively low MIC values of 6.15µM and 1.98µM, both of which are used as antimicrobials in dentistry (8). The estrogen receptor modulating drugs clomiphene citrate, raloxifen, toremifene and tamoxifen citrate (9),(10) displayed liquid MIC values ranging from 9.03µM to 26.64µM (Table 1). GBR12909 (Vanoxerine), a dopamine transport inhibitor (11), displayed a liquid MIC of 26.48 µM (Table 1). Two of the drugs tested against *M. smegmatis*, auranofin and ebselen, displayed MIC values of 0.27µM and 18.5µM respectively while other drugs which were initially identified as hits from the primary screen (fendiline hydrochloride, sulocitidil, apomorphine, nisoldipine, sertraline and fluspirilene) displayed relatively high MIC values that ranged from 77µM to 827.5 µM (Table 1). Some drugs initially examined were excluded from further solid media MIC testing (alexidine dihydrochloride, ebselen and fluspirilene). Fluspirelene displayed a relatively high liquid MIC and alexidine dihydrochloride was discounted for further study due to its structural and functional similarity to chlorhexidine. Further investigation of ebselen ceased due to mode of action deconvolution that has been previously determined elsewhere (12). Ebselen is an organoselenium compound approved by the FDA with a well-known pharmacological profile and is currently being investigated for clinical use in the treatment of bipolar disorders and strokes. Previous studies have shown that ebselen displays antimycobacterial properties and is also effective against multidrug resistant *Staphylococcus aureus* (MRSA) (2). In *M. tuberculosis*, ebselen acts by covalently binding to an active site cysteine residue in antigen 85. Antigen 85 is a complex of secreted proteins (Ag85A, Ag85B and Ag85C) which play an important role in the synthesis of trehalose dimycolates (TDM) and mycolylarabinogalactan (mAG) (12).

**Table 1.**
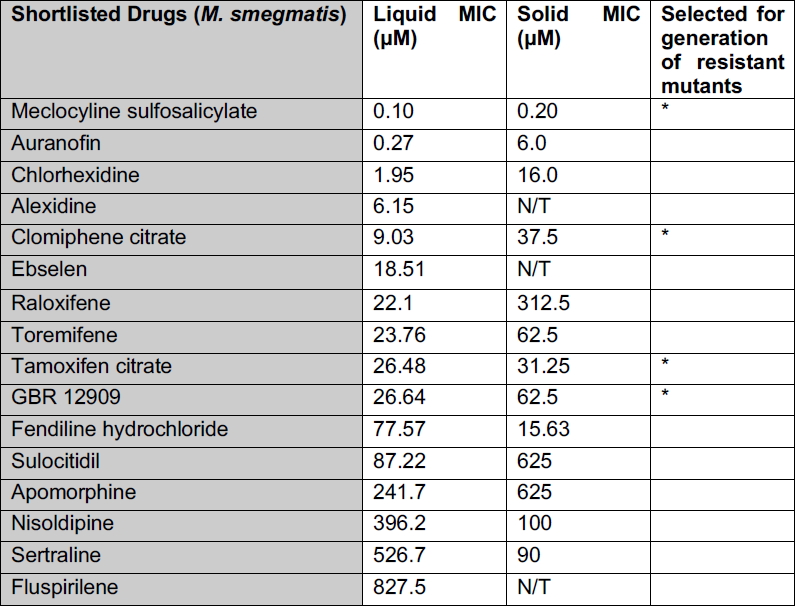
**MIC determination of selected drugs shortlisted as hits from the whole cell screen of the Prestwick Chemical Library ^®^ against** ***M. smegmatis***.MICs were determined using both liquid and solid growth mediums. (N/T) − not tested. (*) – selected for generation of drug resistant mutants.

### Secondary screening and hit confirmation against BCG

MIC studies of the compounds listed in Table 2 revealed thonzonium bromide as having the lowest MIC value of 0.16µM. Thonzonium bromide is a quaternary ammonium monocationic compound which is used as a surfactant and a detergent and has been known to disrupt ATP dependant proton transport in vacuolar membranes along with alexidine dihydrochloride, which are responsible for pH regulation in yeast and *Candida albicans* causing growth defects (13). Florfenicol, a fluorinated analogue of thiamphenicol with broad spectrum activity against Gram negative bacteria and strains resistant to chloramphenicol and thiamphenicol (14), displayed broth and solid MIC values of 0.67µM and 6 µM against *M. bovis* BCG, respectively (Table 2). Florfenicol is known to influence the microbiota of the intestine reducing the amount of uncultured bacterial species similar to *Corynebacterium* and *Mycobacterium* (15). Josamycin, a 16-membered macrolide with inhibitory activity against both Gram negative and Gram positive bacteria (16), displayed potent activity against BCG with an MIC of 0.1µM (Table 2). Interestingly, we identified three antihistamines as having inhibitory activity against BCG. Astemizole had the lowest MIC value of 17.8µM within this group followed by tripelennamine, with an MIC of 41.9µM and olopatadine with the highest MIC value of 202.3 µM in (Table 2). Astemizole (used for general allergies, asthma and rhinitis), tripelennamine (hay fever and rhinitis) and olopatadine (allergic conjunctivitis) are mildly anti-cholinergic and act as H1 receptor antagonists (17–20). Of the antidiabetic drugs displaying activity towards BCG, glipizide and rosiglitazone have an MIC value of 191.1 µM and 43.15 µM, respectively (Table 2). Glipizide is a second-generation sulfonylurea drug that is prescribed for hypoglycaemia in type II diabetes and is known to act by stimulating insulin production and correcting cellular lesions which occur during diabetes mellitus (21, 22). Rosiglitazone, on the other hand, functions by activating peroxisome proliferator activated receptors in adipocytes and sensitising them to insulin (23). Pinaverium, that inhibits L-type calcium channels arresting influx of the Ca^2+^(24), had an MIC of 28.3µM. Two other drugs which displayed relatively high MIC values were granisetron and phentermine (Table 2). Granisetron, an antiemetic drug which is an agonist to the 5-hydroxytryptamine-3 receptor, stimulates the vagus nerve responsible for reflex motility response (25), had an MIC value of 210.6µM. Phentermine, which has been prescribed as an appetite suppressant to control obesity and acts as an agonist to the human TAAR1 (Trace Amine Associate Receptor 1) (26), displayed an MIC of 375.00 µM and was not tested further due to the high concentrations required for inhibitory activity. These drugs were then further tested to establish MICs on solid media in order to determine accurate concentrations to generate spontaneous resistant mutants for mode of action studies. We observed that, upon solid agar MIC testing against BCG, the general trend was that the drugs displayed 5 to 100-fold higher MIC values when compared to MIC values obtained by broth dilution method. For some of the drugs this was attributed to low solubility in solid media as many precipitated during the cooling of the agar medium. Glipizide, olopatadine and granisetron yielded solid MIC values greater than 0.5mM; these drugs precipitated out at higher concentrations and appeared to show no noticeable inhibitory activity against BCG (Fig. S3). Rosiglitazone displayed the highest MIC value at approximately 1.5 mM. Thonzonium and florfenicol had a 5-fold increase in their solid MIC values but were still around 5µM and effectively inhibited the growth of BCG on solid agar (Table 2). Pentamidine, astemizole and pinaverium had solid MIC values of around 0.05 mM while there was a 2-fold increase in the MIC value for tripelennamine (compared to its liquid MIC) of 0.1 mM (Table 2). All compounds listed in Table 1 and Table 2 were tested in an Alamar Blur ^®^ assay against *M. tuberculosis* (Supplementary Information) and the drugs that displayed any notable anti-TB activity are listed in Table 3. Both ebselen and auranofin displayed MICs of 18.51 µM and 0.27 µM, which are in close agreement with previously published values (12, 27). In addition, the estrogen receptor modulating drugs clomiphene and raloxifene inhibited the growth of *M. tuberculosis* H37Rv with MICs of 7.59 µM and 22.10 µM, respectively (we were unable to accurately determine the MIC for Tamoxifen) (Table 3). Finally, GBR12909, used in the clinic to treat cocaine addiction) inhibited the growth of *M. tuberculosis* with an MIC of 26.64 µM. It is interesting to note however, that all of the drugs listed in Table 3 were identified as hits from screens conducted against *M. smegmatis* and not *M. bovis* BCG (Table 1 and Table 2).

**Table 2.** **MIC determination of selected drugs shortlisted as hits from the whole cell screen of the Prestwick Chemical Library ^®^ against ***M. bovis*** BCG.** MICs were determined using both liquid and solid growth mediums. (N/T) − not tested. (*) – selected for generation of drug resistant mutants.

| Shortlisted Drugs<br>(*M. bovis* BCG) | Liquid MIC (µM) | Solid MIC (µM) | Selected<br>generation<br>resistant mutants |
| --- | --- | --- | --- |
| Thonzonium | 0.16 | 5 | * |
| Florfenicol | 0.67 | 6 | * |
| Pentamidine | 3.23 | 50 | * |
| Astemizole | 17.78 | 50 |  |
| Pinaverium | 28.29 | 50 |  |
| Josamycin | 0.10 | 100 |  |
| Tripelennamine | 41.9 | 100 | * |
| Rosiglitazone | 43.15 | ≥ 1500 |  |
| Glipizide | 191.10 | ≥ 500 |  |
| Olopatadine | 202.30 | ≥ 500 |  |
| Granisteron | 210.60 | ≥ 500 |  |
| Phenteramine | 375.00 | N/T |  |

**Table 3.** **MIC determination of selected drugs against** ***M. tuberculosis*** **H37Rv**. MICs were determined in liquid growth media using the Alamar Blue ^®^ assay (Fig. S4).

| Shortlisted Drugs<br>(*M. tuberculosis*) | Liquid MIC (µM) |
| --- | --- |
| Ebselen | 18.51 |
| Clomiphene | 7.59 |
| GBR 12909 | 26.64 |
| Raloxifen | 22.10 |
| Tamoxifen | ≥ 100 |
| Auranofin | 0.27 |

### Generation of spontaneous resistant mutants to determine mode of action

We attempted to generate spontaneous resistant mutants in both *M. smegmatis* and BCG against a selection of drugs identified in Table 1 and Table 2, respectively. However, we were only able to obtain drug-resistant isolates for meclocyline sulfosalicylate, tamoxifen citrate and GBR12909 in *M. smegmatis* (Table 4) and florfenicol, pentamidine and tripelennamine in BCG (Table 5). Analysis of the *M. smegmatis* meclocyline sulfosalicylate mutant revealed a synonymous single nucleotide polymorphism (SNPs) mutation in the gene MSMEG_3619 (*Mtb* ortholog Rv1856c), a probable oxidoreductase deemed non-essential by the Himar-I based transposon mutagenesis ^21^, but also showing importance as having a growth advantage which results in an improvement in fitness when disrupted (28). A single SNP (P122S) was also observed in MSMEG_5249 (*Mtb* ortholog Rv1093) *glyA1* which is a serine hydroxymethyltransferase with possible roles of glycine to serine inter- conversion and the generation of 5, 10-methyenetetrahydrofolate which plays an important role in providing precursors for cellular redox balancing, methylation reactions and a role in thymidylate biosynthesis (Table 4). *GlyA1* is also thought to be an essential gene (28, 29) and it has also been identified as one of the proteins which undergoes PUPylation (Ubquitinylation by prokaryotic ubiquitin protein) in mycobacteria (30). The *M. smegmatis* tamoxifen citrate resistant mutant exhibited a frame shift mutation in the gene MSMEG_6431 (*Mtb* ortholog Rv3849) *espR* which encodes for a protein involved in transcriptional regulation of the three genes Rv3136c-Rv3614c required for the ESX-1 system (Table 4). EspR binds to the promotor region and regulates ESX- 1, therefore controlling virulence of mycobacteria (31). Spontaneous resistant mutants raised against GBR12909 in *M. smegmatis* produced consistent multiple mutations in MSMEG_3033 (*aroB*) (Table 4). *AroB* is predicted to be an essential gene as studied *in M. tuberculosis* (28, 29) and it encodes for 3- dehydroquinote synthase, which is one of several enzymes participating in the shikimate biosynthetic pathway (32). AroB is a homomeric enzyme, is the second enzyme in the shikimate biosynthetic pathway and is present in various bacterial species such as *Corynebacterium glutamicum*, *Escherichia coli*, *Bacillus subtilis* and other fungi, plant and apicomplexan parasites (33–35). AroB makes for an important target due to its essentiality in *M. tuberculosis* and absence of this biochemical pathway in mammals (28, 29, 33). Spontaneous resistant mutants were generated for florfenicol in BCG, revealing a point mutation in BCG_1533 (*echA12*) gene which encodes for a putative enoyl CoA hydratase (Table 5). EchA12 has been shown to be membrane localized within the mycobacterial cell membrane (36), however the gene was not found to be essential through Himar-I based transposon mutagenesis (28, 29). It has been suggested that EchA12 is involved in lipid membrane metabolism and is found to co-localise with thioredoxine A (36) and CtpD, which is an ATPase involved with the metalation of proteins secreted during redox stress (37). Three additional point mutations were also observed in the florfenicol resistant mutant. A single guanine to adenine point mutation in the gene BCG_3185 (PPE50) which encodes for a protein belonging to the PPE family, generates a G251D mutation. BCG_3508 (*rpsI*) encodes for a probable 30S ribosomal protein and contains a P17A mutation in the florfenicol mutant (Table 5). Florfenicol is a fluorinated form of thiamphenicol which belongs to the amphenicol family of antibiotics, whose mode of action is through binding to the 23S rRNA of the 50S ribosomal unit (38). Finally, we also observed a V271A point mutation in BCG_3755c, which encodes for a glycerol kinase (GlpK), which catalyses the rate limiting step in glycerol metabolism of converting glycerol to glycerol-3- phosphate (39, 40). Mutations in *glpK*, have previously been observed when generating resistant mutants to drugs in an attempt to deconvolute their mode of action (41, 42). In our investigation of pentamidine activity, we identified an identical mutation in *glpK*, two separate non-synonymous SNPs in BCG_0763, which encodes for putative membrane protein with a domain of unknown function, and a single SNP in BCG_1609, which encodes for *mmpl6* (Table 5). Mycobacterial membrane protein large (MmpL) are membrane proteins involved in shuttling lipid components across the plasma membrane and have been known to play an important role in drug resistance mechanisms, membrane physiology and virulence of the bacterium (43). The tripelennamine mutant had a single point mutation in the promoter region of BCG_3090, a multi-drug transport integral membrane protein (*mmr*) (Table 5), which is a known efflux pump involved in drug resistance with high susceptibility to quaternary compounds (44). This suggest that exposure to tripelennamine might induce a mutation that causes increased overexpression of Mmr which could alter the ability of the bacterium to efflux drugs.

**Table 4:** **Mode of action determination of drugs inhibiting ***M. smegmatis*** through whole genome sequencing and variant analysis of spontaneous resistant mutants.** The table represents the single nucleotide polymorphisms obtained through whole genome sequencing of the spontaneous resistant mutants raised compared against drug sensitive *M. smegmatis*.

| Drug Name (*M. smegmatis* hits) | Mutated Genes | Rv Number | Positions | Amino acid substitutions | Probable function |
| --- | --- | --- | --- | --- | --- |
| **Meclocycline** | MSMEG_3619 | Rv1856c | A/G |  | Short chain dehydrogenase/ oxidoreductase |
| **Meclocycline** | MSMEG_5249 (*glyA1*) | Rv1093 | Ccg/Tcg | P122S | Serine hydroxymethyltransferase |
| **Tamoxifen** | MSMEG_6431 (*espR*) | Rv3849 | ttc/ (Frame shift) | F24 | Conserved hypothetical protein |
| **GBR12909** | MSMEG_3033 (*aroB*) | Rv2538c | Ggg/Agg | G282R | Involved at the second step in the biosynthesis of chorismate within the biosynthesis of aromatic amino acids (the shikimate pathway) [catalytic activity: 7-phospho-3-deoxy-arabino-heptulosonate = 3-dehydroquinate + orthophosphate]. |
| **GBR12909** | MSMEG_3033 (*aroB*) | Rv2538c | gGc/gAc | G284D |  |
| **GBR12909** | MSMEG_3033 (*aroB*) | Rv2538c | Tgc.GTtgc (Frame shift) | C356V |  |
| **GBR12909** | MSMEG_3033 (*aroB*) | Rv2538c | cTa/cCa | L363P |  |

**Table 5:** Mode of action determination of drugs inhibiting ***M. bovis BCG*** through whole genome sequencing and variant analysis of spontaneous resistant mutants. The table represents the single nucleotide polymorphisms obtained through whole genome sequencing of the spontaneous resistant mutants raised compared against drug sensitive *M. bovis* BCG. Drug Name

| Drug Name<br>(*M. bovis* BCG hits) | Mutated Genes | Rv Number | Positions | Amino acid substitutions | Probable function |
| --- | --- | --- | --- | --- | --- |
| **Florfenicol** | BCG_1533<br>(EchA12) | Rv1472 | Gga/Aga | G239R | Possible enoyl-CoA hydratase echA12 [*Mycobacterium bovis* BCG str. Pasteur 1173P2] |
| **Florfenicol** | BCG_3158<br>(PPE50) | Rv3135 | gGc/gAc | G251D | PPE family protein |
| **Florfenicol** | BCG_3508<br>(rpsI) | Rv3442c | Ccc/Gcc | P17A | Probable 30S ribosomal protein S9 RPSI |
| **Florfenicol** | BCG_3755c<br>(glpK) | Rv3696c | gTc/gCc | V271A | Probable glycerol kinase GlpK (ATP glycerol 3-phosphotransferase) |
| **Pentamidine** | BCG_0763 | Rv0713 | aTt/aCt | I274T | Probable conserved transmembrane protein [*Mycobacterium bovis* BCG str. Pasteur 1173P2] |
|  | BCG_0763 | Rv0713 | gCg/gTg | A281V | Probable conserved transmembrane protein [*Mycobacterium bovis* BCG str. Pasteur 1173P2] |
| **Pentamidine** | BCG_1609<br>(mmpL6) | Rv1557 | Gcc/Acc | A31T | Probable conserved transmembrane protein |
| **Pentamidine** | BCG_3755c<br>(glpK) | Rv3696c | gTc/gCc | V271A | Probable glycerol kinase GlpK (ATP glycerol 3-phosphotransferase) |
| **Trippelennamine** | promoter region of gene BCG_3090 | Rv3065 | Upstream gene BCG 3089c<br>(Rv 1904) |  | Multi-drug transport integral membrane protein (efflux pump) |
|  | glpK | Rv3696c | gTc/gCc | V271A | Probable glycerol<br>kinase GlpK (ATP<br>glycerol 3-phosphotransferase) |

## Discussion

Screening compounds against *M. smegmatis* has a distinct advantage over the slower growing *M. bovis* BCG strain in terms of its shorter generation time, thus expediting the generation of screening data and turnaround of results. However, using *M. smegmatis* as a screening organism is less efficient in determining antitubercular compounds than *M. bovis* BCG. It was observed during a screen of the LOPAC library against *M. tuberculosis, M. smegmatis* and *M. bovis* BCG that 50% of the drugs inhibiting *M. tuberculosis* were not identified in *M. smegmatis* while it was only 21% of the drugs that were not identified in *M. bovis* BCG. In addition, it was observed that 30% of proteins in Mtb do not have conserved orthologs in *M. smegmatis* (5). Despite this fact, bedaquiline, the most recent drug given FDA approval for the treatment of MDR-TB, was initially discovered through a whole cell screen assay against *M. smegmatis* (45), which makes the case for not excluding *M. smegmatis* as a model organisms for antitubercular drug screening.

Although genetically similar, there are a number of physiological variations between *M. tuberculosis* and *M. bovis* BCG which have been attributed to the differential expression of around 6% of genes across their respective genomes. During the exponential growth of both organisms, major variations were observed for genes involved in cell wall processes, intermediary metabolism and respiration and hypothetical proteins (46). In addition, in *M. tuberculosis* the PE/PPE genes were found to be highly expressed whereas in *M. bovis* BCG, there are a higher number of transcriptional regulators that are overexpressed during exponential growth. These variations in gene expression profiles of mycobacteria partly explain why different classes of drugs differentially inhibit each strain utilised during screening experiments (Fig. 4).

Target deconvolution of hits that emerge from whole cell screening efforts has long been the bottleneck of phenotypic-based drug discovery; often huge investment of time and resource is required to identify the precise molecular target of active compounds (47). Interestingly, in the case of early stage drug discovery in *M. tuberculosis*, there seems to be an apparent trend whereby inhibitors of cell growth/viability obtained through phenotypic screening efforts tend to inhibit membrane targets such as DprE1, MmpL3, QcrB and Pks13 (48). This relatively high probability of hits inhibiting membrane targets could, in part, be due to the hydrophobicity of the inhibitors screened in libraries against mycobacteria (48, 49). For instance, several drugs from the Prestwick Chemical Library ^®^ that inhibit the growth of mycobacteria have an average clogP value of 5.7 (48). Hydrophobic drugs have a tendency to enter into the lipid layers of the mycobacterial cell envelope and then move laterally through the membrane due to their inability to cross the plasma membrane into the cytoplasm. While traversing the bilayer, these hydrophobic compounds interact with membrane proteins and thus there is an increased probability of the drugs inhibiting such targets thereby producing membrane protein related mutations in spontaneously generated mutants during mode of action studies (48). Alexidine dihydrochloride and thonzonium bromide were amongst the hits observed in this study that have uncoupling properties and might generated a membrane protein mutation (13). In this study, screening the Prestwick Chemical Library ^®^ also identified inhibitors such as calcium channel blockers, antihistamines, antifungal azoles and, unsurprisingly, a variety of antinfectives (Fig. 4). Calcium channel inhibitors are generally small hydrophobic molecules which have the ability to enter the phospholipid bilayer and can diffuse through the membrane inhibiting metabolic functions due to interactions with proteins and boundary lipids (50). Antifungal azoles, which have been shown to elicit inhibitory activity against mycobacteria, act by targeting the CYP121 and CYP130 cytochrome P450 systems (51). Our screening of the Prestwick Chemical Library ^®^ and analysis of unique emerging hits, provides new impetus to explore drug repurposing as a feasible and efficient way of mining for new anti-mycobacterial drugs.

## Materials and Methods

### Bacterial strains, plasmids and growth media

*M. smegmatis* mc^2^155 was electroporated with pSMT3-eGFP and transformants were selected on Tryptic Soy Agar supplemented with hygromycin B (20 µg/ml). Single colonies were used to inoculate 10 mL of Tryptic Soy Broth supplemented with Tween 80 (0.05% v/v) at 37°C with shaking at 180 rpm. *M. smegmatis* mc^2^155 harbouring pSMT3-eGFP was diluted 1/100 into Middlebrook 7H9 supplemented with glycerol (2 mL/L) and Tween 80 (0.05% v/v) and further sub-cultured at 37°C with shaking at 180 rpm.*M. bovisn* BCG was electroporated with pSMT3-eGFP and transformants selected on Middlebrook 7H10 containing OADC (10% v/v) and hygromycin B (20 µg/ml). Single colonies were inoculated into 50 mL of Middlebrook 7H9 containing OADC (10% v/v) and Tween 80 (0.05% v/v) and statically cultured at 37°C for ˜ 5 days. Both *M. smegmatis* mc^2^155 and *M. bovis* BCG expressing eGFP were quantified by sampling 200 µL of cells which were 2-fold serially diluted across a black F-bottom 96-well micro-titre plate and fluorescence was measured using a BMG Labtech POLARstar Omega plate reader (Excitation 485-12 nm, Emission 520 nm).

### Validation of eGFP reporter screen

Batch cultures of *M. smegmatis* pSMT3-eGFP and *M. bovis* BCG pSMT3-eGFP were adjusted to give a basal reading of 20,000 Relative Fluorescent Units (RFU) by diluting into fresh Middlebrook 7H9 containing Tween 80 (0.05% v/v) with additional OADC (10% v/v) for *M. bovis* BCG (final well volume of 200 µL). The anti-mycobacterial drugs isoniazid, ethambutol, streptomycin, pyrazinamide and rifampicin were included in over a range of concentrations on assay plates. Wells containing mycobacterial culture in the presence of 1 % DMSO represent high controls, whilst wells containing only media constitute low controls. Assay plates were cultured for 48 hours in a ThermoCytomat plate-shaker incubator (100 % humidity at 37°C with 180 rpm plate agitation) and eGFP fluorescence was measured kinetically every 2 hours.

### Medium Throughput Screen of the Prestwick Chemical Library ^®^

The Prestwick Chemical Library ^®^ (1200 drugs) was purchased from Specs.net preformatted in master plates so that all compounds were solubilized in 100 % DMSO at a final concentration of 10 mM. A fully automated Hamilton Star workstation was used for all liquid handling protocols. Compounds were loaded into black F-bottom 96-well assay ready plates (Greiner) followed by 200 µL of either *M. smegmatis* pSMT3-eGFP or *M. bovis* BCG pSMT3-eGFP resulting in a final drug concentration of 20 µM in the primary screen. Wells containing only cells (high control) or cells in combination with 50 µg/mL rifampicin (low control) were included on each assay plate to establish positive and negative controls, respectively. Assay plates were cultured at for 48 hours in a Thermo Cytomat plate-shaker incubator (100 % humidity at 37°C with 180 rpm plate agitation) and eGFP fluorescence was measured kinetically every 2 hours to generate growth curves for individual wells of each assay plate. Data from the final 48 hr read was normalized using the following equation:

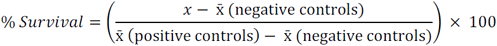

Each assay plate was checked for robustness and reproducibility by calculating the Z’-factor using the following equation;

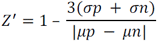

For the primary screen, positive controls and negative controls were included in columns 1 and 2 respectively. The Z’ was found to be on average 0.75, well above the Z’>0.5 which is widely regarded as being suitable for HTS. All 1200 drugs from the Prestwick Chemical Library ^®^ were screened in duplicate and hits were identified as inhibiting cell growth by ≥75%, as determined by measuring eGFP fluorescence.

### Validation of selected hits and MIC determination in liquid media

Drugs selected for further study were purchased from a variety of commercial vendors. Drugs were dissolved into 100% DMSO resulting in a 10 mM stock that was subsequently used to generate a 10-point 3-fold serial dilution which provided a dose response curve with maximum and minimum drug concentration of 500 µM and 0.0254nM, respectively. Data was normalised as described above. The concentration of drug that is required to inhibit cell growth by 99% was calculated by non-linear regression (Gompertz equation for MIC determination, GraphPad Prism).

### MIC determination on solid agar

Selected compounds identified from the secondary MIC screen were further tested for MIC evaluation using solid agar media. Drugs were diluted from a 10mM stock, mixed individually in 2mLs of molten 7H10 agar and dispensed in square partitioned petri plates. Plates were incubated at 37°C and solid MIC was determined based on absence of colonies.

### Spontaneous mutant generation

Spontaneous mutants were generated by plating 10^8^ cells (*M. smegmatis* and/or *M. bovis* BCG) per plate on 7H10 agar with drug concentrations of 2.5x, 5x and 10x the solid MIC values. Plates were incubated at 37 °C for a week or 30 days for *M. smegmatis* and *M. bovis* BCG, respectively. Colonies that appeared on plates containing drug at 10X MIC were inoculated into 7H9 re- plated onto 7H9 agar in the presence drugs at 10X MIC in order to confirm resistance. Genomic DNA was isolated for both wild type *M. smegmatis* and *M. bovis* BCG strains together with resistant mutants (52, 53). Genomic DNA was submitted to MicrobesNG (https://microbesng.uk/) for whole genome sequencing and SNP variant analysis of the sequence in comparison to the wild type genomic DNA for each strain.

## Supplemental Material

Supplementary material (Figures S1 to S4) for this article are provided for online publication.

